# Chromosome-level genome assembly of milk thistle (*Silybum marianum* (L.) Gaertn.)

**DOI:** 10.1101/2024.02.29.582772

**Authors:** Kyung Do Kim, Jeehyoung Shim, Ji-Hun Hwang, Daegwan Kim, Moaine El baidouri, Soyeon Park, Jiyong Song, Yeisoo Yu, Keunpyo Lee, Byoung-Ohg Ahn, Su Young Hong, Joong Hyoun Chin

## Abstract

*Silybum marianum* (L.) Gaertn., commonly known as milk thistle, is a medicinal plant belonging to the Asteraceae family. This plant has been recognized for its medicinal properties for over 2,000 years. However, the genome of this plant remains largely undiscovered, having no reference genome at a chromosomal level. Here, we assembled the chromosome-level genome of *S. marianum*, allowing for the annotation of 53,552 genes and the identification of transposable elements comprising 58% of the genome. The genome assembly from this study showed 99.1% completeness as determined by BUSCO assessment, while the previous assembly (ASM154182v1) showed 36.7%. Functional annotation of the predicted genes showed 50,329 genes (94% of total genes) with known protein functions in public databases. Comparative genome analysis among Asteraceae plants revealed a striking conservation of collinearity between *S. marianum* and *C. cardunculus* . The genomic information generated from this study will be a valuable resource for milk thistle breeding and for use by the larger research community.

## Background & Summary

*Silybum marianum* (L.) Gaertn., commonly known as milk thistle, is an annual or biennial plant belonging to the *Asteraceae* family^1–3^ and has been recognized for its medicinal properties for over 2,000 years^4,5^. Silymarin, a complex of flavonolignans extracted from milk thistle seeds^6–8^, exhibits remarkable hepatoprotective and detoxifying effects^9–14^. In recent years, it has garnered attention as a potential therapeutic agent for various liver ailments, including alcoholic liver disease and acute viral hepatitis^15–19^.

Despite being a distinct species from *Cirsium* spp., milk thistle is often misidentified due to phenotypic similarities. Therefore, deciphering the milk thistle genome holds immense value in understanding and optimizing the plant’s beneficial properties. Sequencing the milk thistle genome can help researchers understand the molecular mechanisms underlying silymarin’s therapeutic properties and identify new compounds with potential medicinal applications. Additionally, it can help identify genes that can be manipulated to increase silymarin production. This knowledge can also help develop strategies to protect plants from pests and diseases. Despite the growing recognition of silymarin’s therapeutic potential, the genome of *S. marianum* remains largely uncharted. Yet, there is no reference genome sequenced for *S. marianum* at the chromosomal level. This lack of genomic resources poses a significant hurdle to advancing research and plant breeding on *S. marianum*.

To bridge this gap, we assembled the chromosome-level genome of *S. marianum* using a combination of Oxford Nanopore long-read, Illumina short-read, and Pore-C technologies. This study unveiled the genetic landscape and diversity of the plant, allowing for the annotation of 53,552 genes and the identification of transposable elements consisting of 58% of the genome. The genomic resources, gene structure, and functional insights generated from this study will pave the way for future research efforts aimed at harnessing the full potential of milk thistle.

## Methods

### Sample preparation and genomic sequencing

*Silybum marianum* cv. ‘Silyking’, also known as ‘EM05’, is a patented variety recognized for its abundant silymarin content (Figure 1A). EM05 originated from germplasm collected in 2017 at local farms in Pyeongtaek, Gyeonggi-do, Korea. It was carefully selected from heterogeneously collected accession and self-propagated to achieve the pure line of ‘EM05’ by EL&I Co., ltd. in Hwaseong, Gyeonggi-do, Korea. Genomic DNA was extracted from young leaves of EM05 using the Cetyltrimethylammonium Bromide (CTAB) method. The quality and quantity of the extracted DNA were assessed using NanoDrop 2000 (Thermo Fisher Scientific, USA).

**Figure 1.**
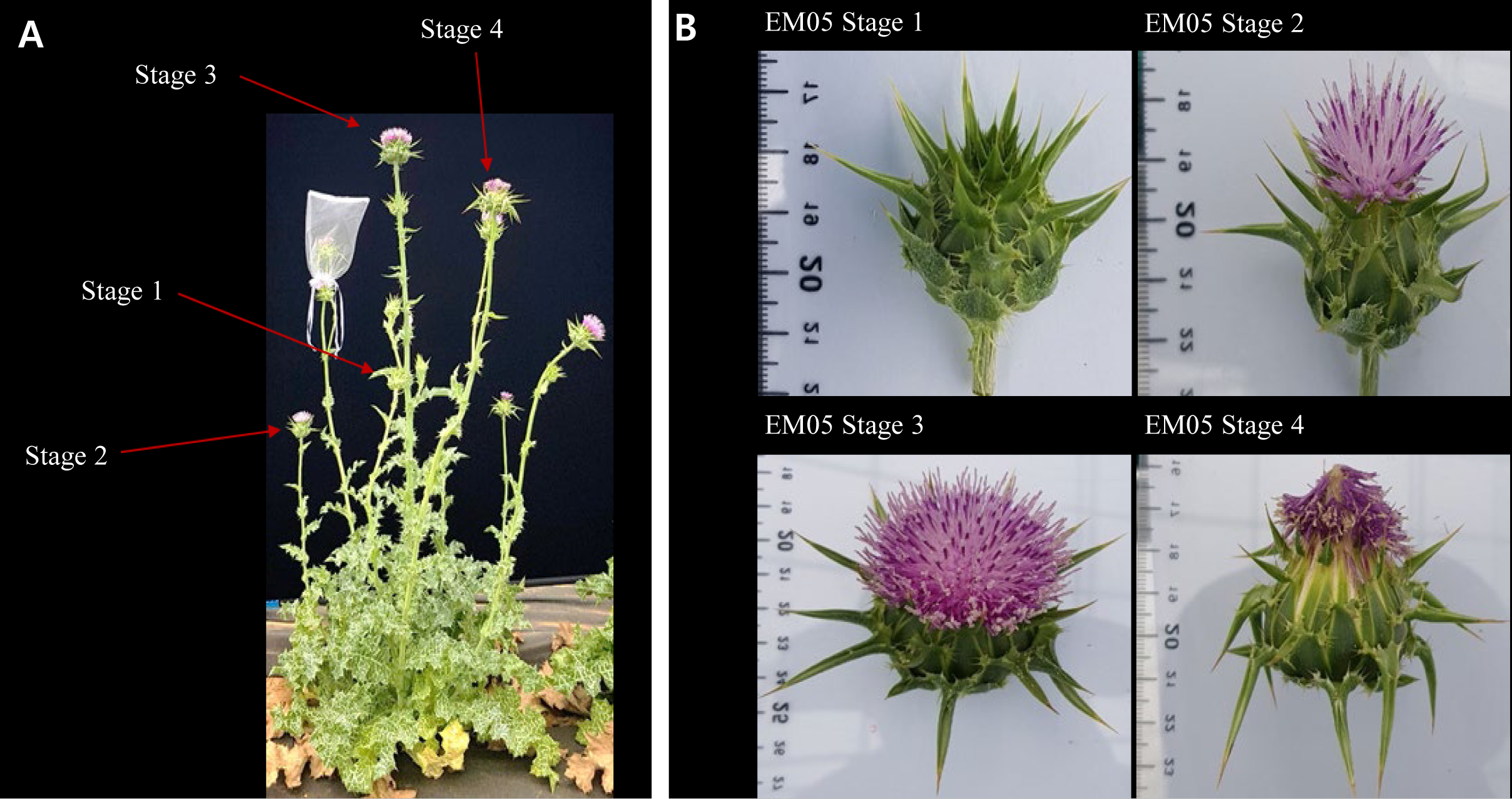
Morphology and flowering stages of *Silybum marianum* . (A) Morphology of *S. marianum* plant. Each arrow reflects different flowering stages. (B) Flowering stages of *S. marianum*. Stage 1: No petals have emerged, and small white seeds are visible near the base of the flower receptacle. Stage 2: Some petals have emerged, and small white seeds are visible near the base of the flower receptacle. Stage 3: Most petals have emerged, but they are not yet withered. Slightly larger white seeds are visible near the flower receptacle. Stage 4: Most petals have emerged, but they are withered. The flower receptacle has thickened, and the seeds are larger and firmer.

Nanopore library was prepared using a ligation sequencing kit, SQK-LSK110. Long-read sequencing was performed using an FLO-PRO002 flow cell on the Oxford Nanopore PromethlON platform. A total of 77.31 Gb raw data with an average read length of 25.24 Kb and an N50 length of 39.84 Kb were obtained, accounting for ∼ 111.3-foldsCof the genome (Table 1). Illumina paired-end library with a 400 bp insert size was prepared using the TruSeq Nano DNA kit. Short-read sequencing was conducted on the Novaseq 6000 platform with 2 × 150 bp reads, which generated 52.24CGb raw data, accounting for ∼ 75.23-foldsCof the genome (Table 1). The low-quality sequences with a Phred score of 20 or lower, as well as Illumina adapter sequences, were removed using Trimmomatic v.0.39^20^.

**Table 1.**
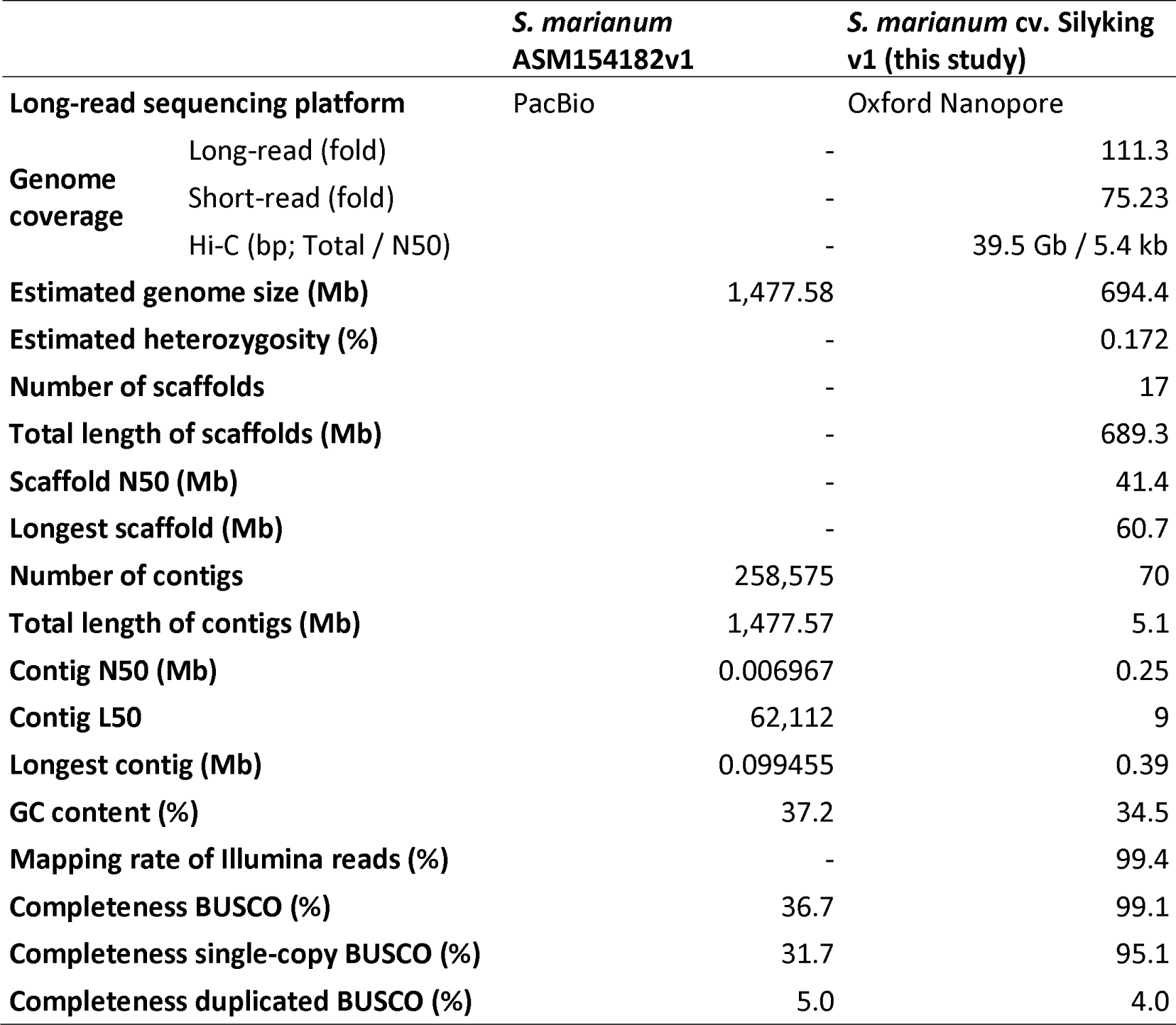
Comparison of genome assemblies between *Silybum marianum* ASM154182v1 and cv. Silyking v1.

### Transcriptome sequencing

For RNA extraction, seven tissue samples from various parts of the *S. marianum* EM05 plant, including flowers, leaves, stems, and roots, were collected. For flower tissues, samples from four different stages of inflorescence were collected (Figure 1B). Total RNA was extracted using CTAB buffer (OPS Diagnostics, USA) with the addition of 50µL of β-Mercaptoethanol to 500ml of the buffer. The process involved mixing the samples with 900µL of CTAB buffer, followed by centrifugation at 14000rpm for 5 minutes at 4C. The resulting 1st supernatant (700µL) was then incubated at 65C for 15 minutes with intermittent vortexing. The lysate was mixed with an equal volume of Phenol: chloroform: isoamyl alcohol (25:24:1) PCI and centrifuged at 14000rpm for 5 minutes at 4C. Subsequently, the 600µL supernatant was mixed with LiCl(5M) in a 1:1 ratio, incubated at -20C for 4 hours, and then centrifuged at 14000rpm for 10 minutes at 4C. After removing the supernatant, a 500µL wash with 70% Ethanol was performed, followed by centrifugation at 14000rpm for 3 minutes at 4C. The samples were air-dried for 20 minutes before adding 50µL elution buffer (0.1x TX buffer) with thorough mixing. For DNase1 treatment, QIAGEN DNase1 powder was dissolved in 550µL H2O and then aliquoted into 1.5ml E-Tube in each tube. Just before use, buffer was added to DNase1 in a 1:1 ratio. Incubation was carried out at 37C for 30 minutes. RNA sequencing library was prepared using TruSeq Stranded mRNA Sample Preparation Kit and sequenced on the Novaseq 6000 platform with 2 × 101 bp reads. A total of 36 Gb of raw data with an average of 52 million reads per sample was generated from seven *S. marianum* samples.

### Genome assembly and chromosome-level scaffolding

The characteristics of the *S. marianum* genome were estimated based on a total of 304,981,656 trimmed Illumina read pairs with 151 bp in length. The distribution of *k*-mer read depth was computed using Jellyfish v2.2.10^21^, and the genome size and heterozygosity were calculated using GenomeScope v2.0^22^ with default parameters. In this study, *k*-mer values of 19 and 21 were used. The estimated genome size was 643 Mb with 0.14% heterozygosity using 19-mer and 654 Mb with 0.14% heterozygosity using 21-mer (Table 1, Supplementary Figure 1).

The draft genome of *S. marianum* was assembled using Oxford Nanopore long-reads with Nextdenovo v2.5.0^23^. The assembly resulted in 70 contigs with a total length of 706 Mbp. Gap sequences in the draft genome were polished using Illumina short-reads with NextPolish v1.4.0^24^.

To assemble the chromosome-level genome, a Pore-C library was prepared. This involved various steps such as nuclei isolation, chromatin denaturation, digestion, ligation, de-crosslinking, and DNA extraction. Library construction was carried out using the extracted DNA and the SQL-LSK110 ligation kit (Oxford Nanopore) following the manufacturer’s protocol. The constructed libraries were checked for quality on a 1.0% TBE agarose gel. The Pore-C library was sequenced using the Oxford Nanopore PromethION platform, generating 39.47 Gbp of raw data (Table 1). The raw data was trimmed using Guppy v3.0.4 with Q>=7, resulting in 34.05 Gbp of raw data with a mean quality of 11.9. Only the trimmed data was statistically assessed with anoPlot v1.40.0. Mapping of trimmed Pore-C data to the assembly and removal of duplicated alignments were performed using Pore-C Snakemake v0.4.0. Assembly, hic, and fastq files were created using 3D-DNA pipeline v180922. The assembly was manually curated based on the pairwise contact heatmap (Figure 2A) generated using JuiceBox v1.11.08^25^. After scaffolding, a total of 35 contigs were connected into 17 chromosome-level scaffolds with a total length of 689.3 Mbp (Supplementary Table 1). Unplaced contigs showing high similarity with bacterial sequences were excluded from the assembly, resulting in the exclusion of 10 contigs with a total length of 6.7 Mbp (Supplementary Table 2).

**Figure 2.**
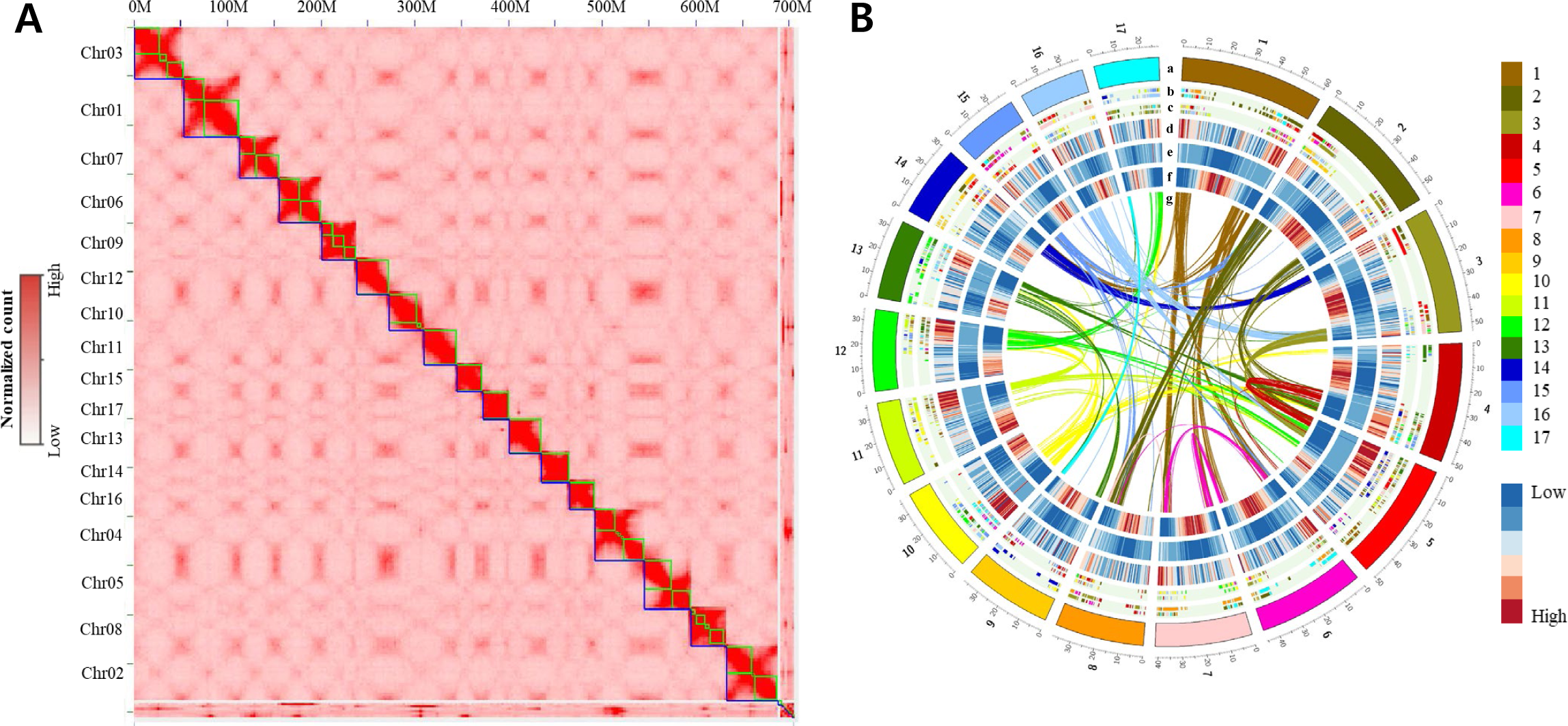
Overview of the genomic landscape of *Silybum marianum.* (A) Pore-C interaction heatmap of *S. marianum* assembly. The interactions of the *S. marianum* chromosome were measured by the number of the Pore-C reads illustrated by red color. (B) Genome features of *S. marianum* across the 17 chromosomes. Each track was drawn in a 500 kb window. The outer to the inner tracks represent: (a) Chromosomes of *S. marianum*; (b) Synteny regions between *Cynara cardunculus* and *S. marianum* ; (c) Synteny regions between *Helianthus annuus* and *S. marianum* ; (d) Gene count of *S. marianum* in 500 kb; (e) DNA TE count of *S. marianum* in 500 kb; (f) LTR TE count of *S. marianum* in 500 kb. (g) Curved lines at the center show segmental duplication regions in *S. marianum* . Each color labeled at the track a, b, c, and g represents each chromosome.

**Figure 3.**
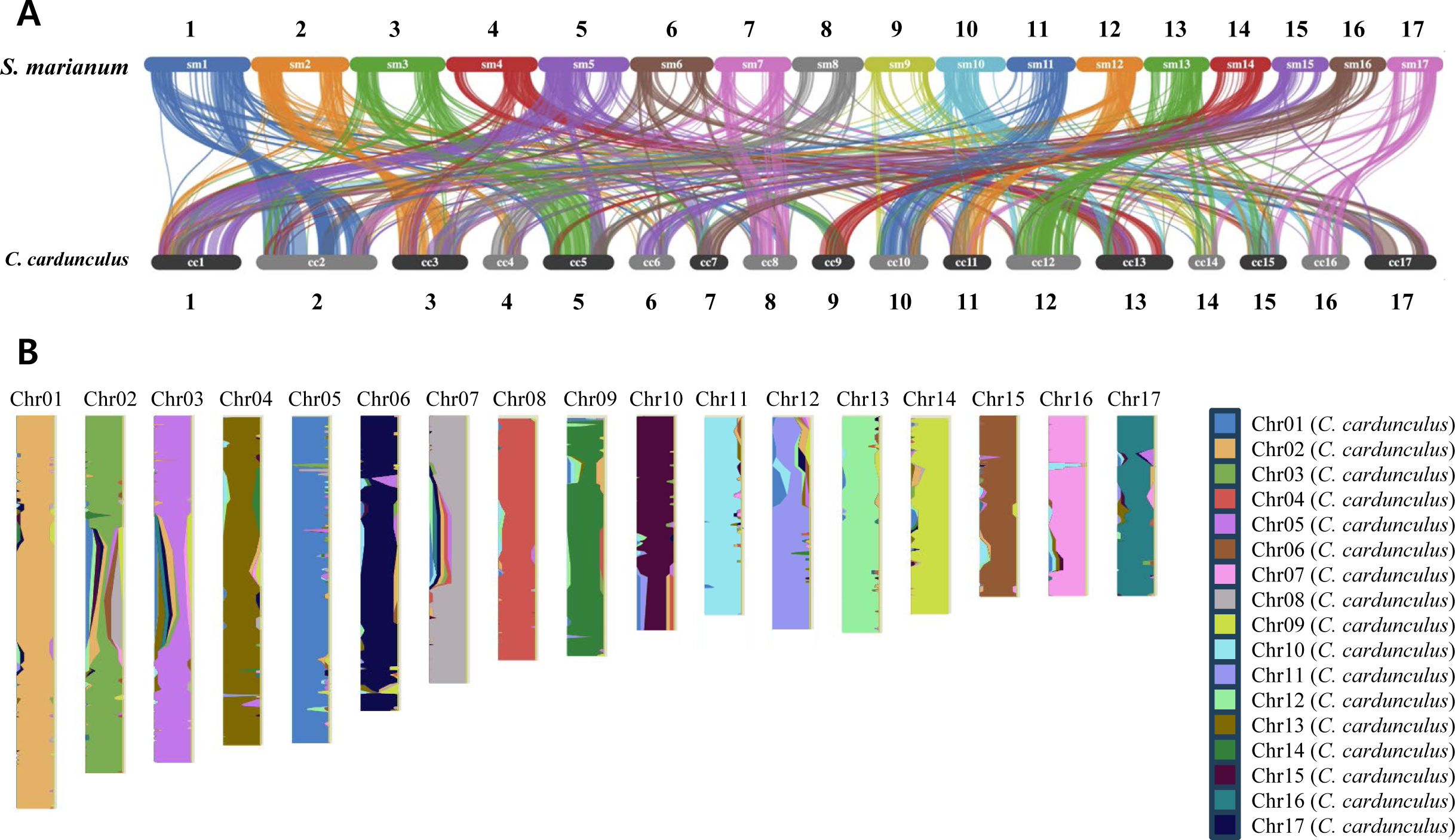
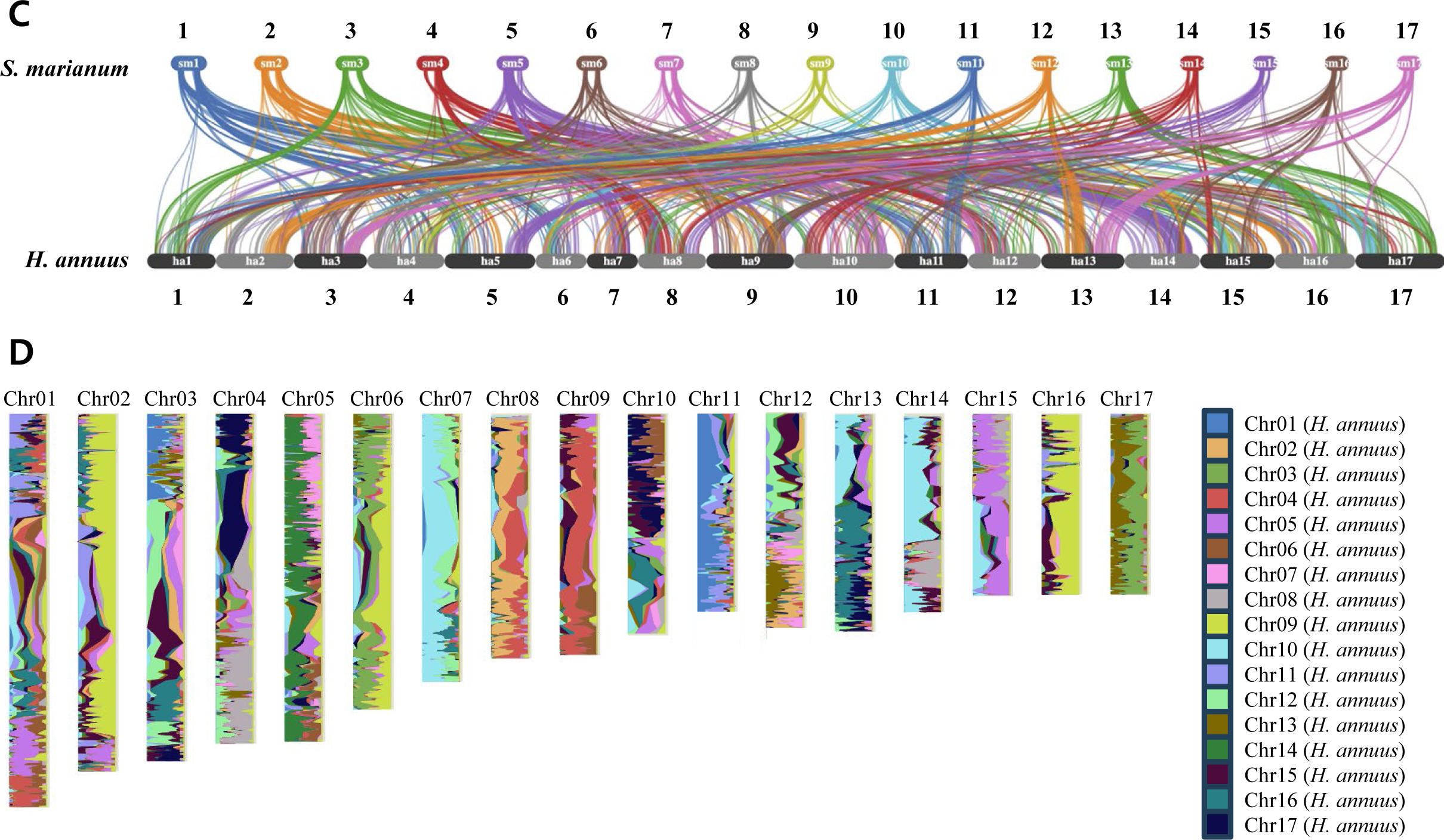
Comparative genome analysis between *Silybum marianum, Cynara cardunculus,* and *Helianthus annuus* . (A) Collinearity between *S. marianum* and *C. cardunculus* and across 17 chromosomes. (B) *S. marianum* chromosomes painted with collinearity regions between *S. marianum* and *C. cardunculus*. (C) Collinearity between *S. marianum* and *H. annuus* across 17 chromosomes. (D) *S. marianum* chromosomes painted with collinearity regions between *S. marianum* and *H. annuus*.

### TE annotation

The annotation of transposable elements (TEs) was conducted via both homology and structural search procedures. The initial step involved aligning multiple TE protein databases, including Repbase (version 19.06), REXdb^26^, and TREP databases, against the *S. marianum* reference genome using the fastx32 program with an e-value of 1e-5. Once the alignment was completed, overlapping genomic intervals for each TE and superfamily were merged utilizing Bedtools merge, taking into consideration the insertion strand (-s option). The corresponding nucleotide sequences were subsequently extracted in Fasta format for each superfamily. An ’all-against-all’ BLASTn search was executed for each superfamily using a minimum e-value of 1e-50. Clustering of different families was performed using the SiLiX program^27^ with a minimum of 80% of identity over 80% of coverage. At this stage of the annotation process, the TE sequences identified represented only the coding regions of the elements, and precise element boundaries were still undefined. Thus, for each paralog within the same family, 10 kbp flanking regions were extracted, and alignment was performed using pblat^28^ to redefine the exact TE boundaries by excising regions that lacked alignment with other paralogs. Once the correct boundaries were identified, multiple sequence alignments were performed using MAFFT^29^, and consensus sequences were generated. This resulted in a total of 408 Class I and 129 Class II elements with consensus sequences. Additionally, we ran LTRharvest^30^ using default parameters except for -xdrop 37 - motif tgca -motifmis 1 -minlenltr 100 -maxlenltr 3000 -mintsd 2. Similar to the strategy described earlier, paralogs were then clustered using SiLiX ^27^, and consensus sequences for each family were generated. In total, we identified 563 long terminal repeat (LTR) families. Miniature inverted-repeat transposable elements (MITEs) were identified using MITE-tracker with default parameters. This resulted in the characterization of 443 non-redundant families. TEs identified using in-house strategy, LTR_harvest, and MITE-tracker were merged and redundant families removed, which gave rise at the end to 1239 consensus TE sequences, including 270 Gypsy LTRs, 265 Copia LTRs, 17 LINEs, 49 Mutator, 25 CACTA, 10 Harbinger, 19 Helitrons, and 443 MITEs.

Using the newly characterized 1239 consensus TE sequences, we found that TEs make up 58% of the *S. marianum* genome (Table 2). Most of these elements were located in the pericentromeric regions of chromosomes (Figure 2B, Supplemental Figure 2). In comparison with other plant genomes, a similar pattern was observed where LTRs emerged as the predominant TE type, contributing 59% of total TEs in this species. In Class II, MITEs emerged as the most abundant among terminal inverted repeat transposons (TIRs), accounting for 11.2 % of the total genome.

**Table 2.**
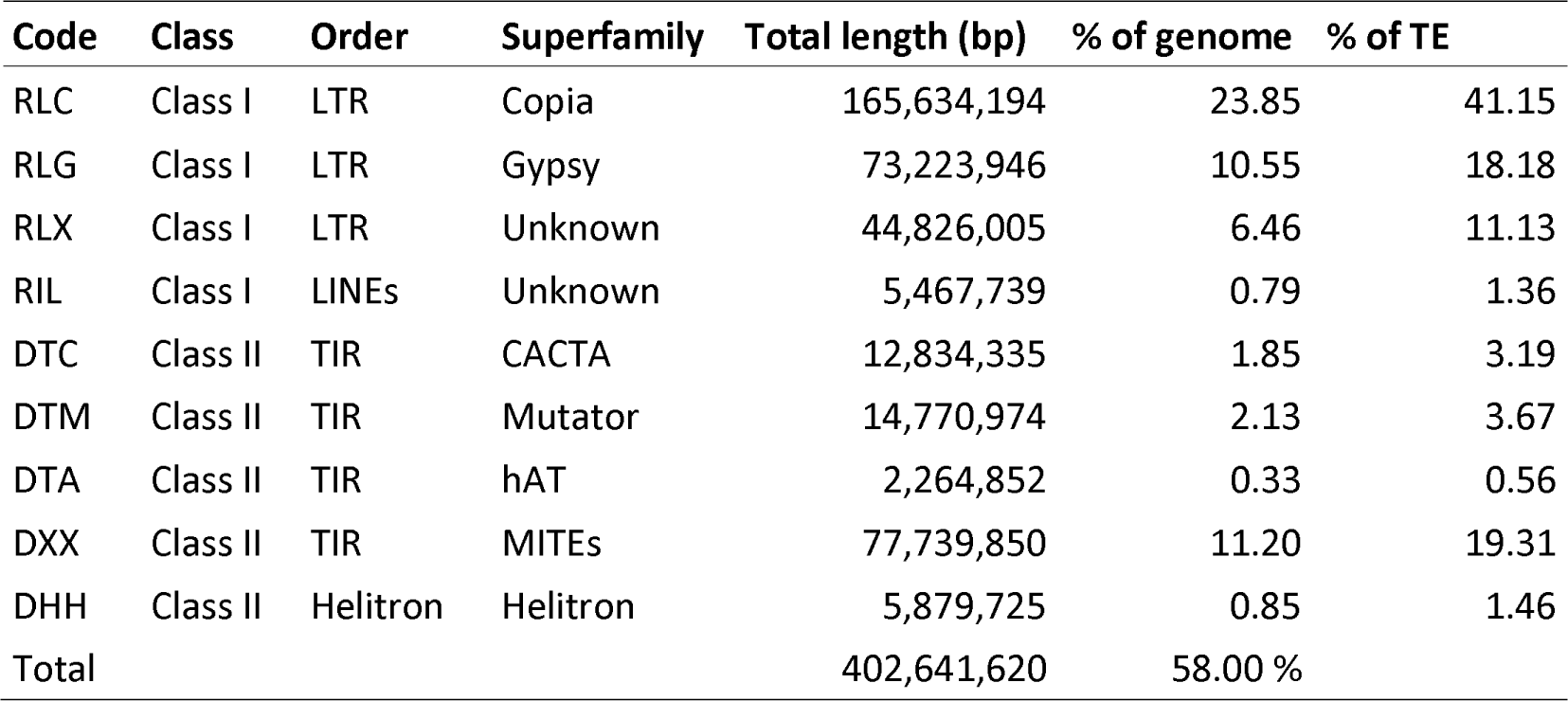
Summary of transposable elements in *Silybum marianum* cv. Silyking v1. Code Class Order Superfamily Total length (bp) % of genome % of TE.

### Gene prediction and functional annotation

Protein-coding genes in the assembled genome were predicted using a combination of *ab initio* prediction, transcriptome-based prediction, and protein alignment. Repetitive sequences in the *S. marianum* genome were masked using RepeatMasker v4.0.5. Raw sequences of RNA-seq data were pre-processed (trim, filter, and remove adapters) using Trimmomatic v0.39^20^ with a Q>20 and 50 bp minimum read length threshold. High-quality reads were then aligned to the assembly using HISAT2 v2.1.0^31^, achieving an average alignment rate of 97.6%. The *ab initio* prediction was carried out with the assistance of BRAKER v1.11^32^, GeneMark-ES/ET v4.48-3.60^33,34^, and AUGUSTUS v3.2.2^35^, utilizing the mapped RNA-seq reads and the assembly with repeat sequences masked. This approach predicted 192,663 genes with a mean exon length of 384 bp. For the transcriptome-based prediction, the high-quality RNA-seq reads were assembled *de novo* using Trinity v2.8.6^36^. The RNA-seq reads were then mapped to the transcriptome assembly and annotated using StringTie v2.0.4^37^. The *de novo* transcriptome assembly and mapped read annotation were aligned against the genome assembly to model complete and partial gene structures using PASA v2.4.1^38^, resulting in the prediction of 101,524 genes with a mean exon length of 321 bp. In addition, the evidence-based gene models were generated using Exonerate v2.2.0^39^ based on the protein sequences of closely related species of *S. marianum* . This approach predicted 52,185 genes with a mean exon length of 250 bp. Lastly, the gene prediction models from *ab initio* prediction, transcriptome-based prediction, and protein alignment were integrated using EvidenceModeler v1.1.1^40^ with different weightings assigned. Subsequently, coding genes lacking start or stop codons or originating from transposable elements were excluded using BLAST v2.9.0, resulting in the prediction of a total of 133,358 gene models.

To investigate the functions of the 133,358 gene models, BLASTp v2.9.0 search was conducted against NCBI plant Refseq DB (7,734,553 sequences), Uniprot DB (565,254 sequences), and TAIR DB (48,356 sequences). In addition, conserved protein domain, gene ontology, and pathway analyses necessary for gene function inference were performed based on Pfam, GO, and KEGG databases using InterProScan v5.38^41^. A gene was considered expressed if the read count within the integrated gene model region in the RNA-seq alignment exceeded zero. The results of the BLASTp, InterProscan, and RNA-seq alignment analyses revealed that 79,862 of the gene models had either associated function or transcript evidence, while 5,779 genes curated as polyproteins were excluded. As a result, 74,083 (55.55%) gene models were selected. Subsequently, a total of 36,163 genes that overlapped with transcriptome-based prediction or protein alignment results were selected. Additionally, 21,266 genes that did not overlap with transcriptome-based prediction or protein alignment but had descriptions at BLASTp and InterProScan results were selected. A total of 3,447 genes with hits to the bacterial genome and 430 genes without hits to the *S. marianum* assembly were excluded. Lastly, a total of 53,552 genes were selected as final gene models with a mean exon length of 289 bp and an average of 3.9 exons per gene (Table 3, Supplementary Table 3).

**Table 3.**
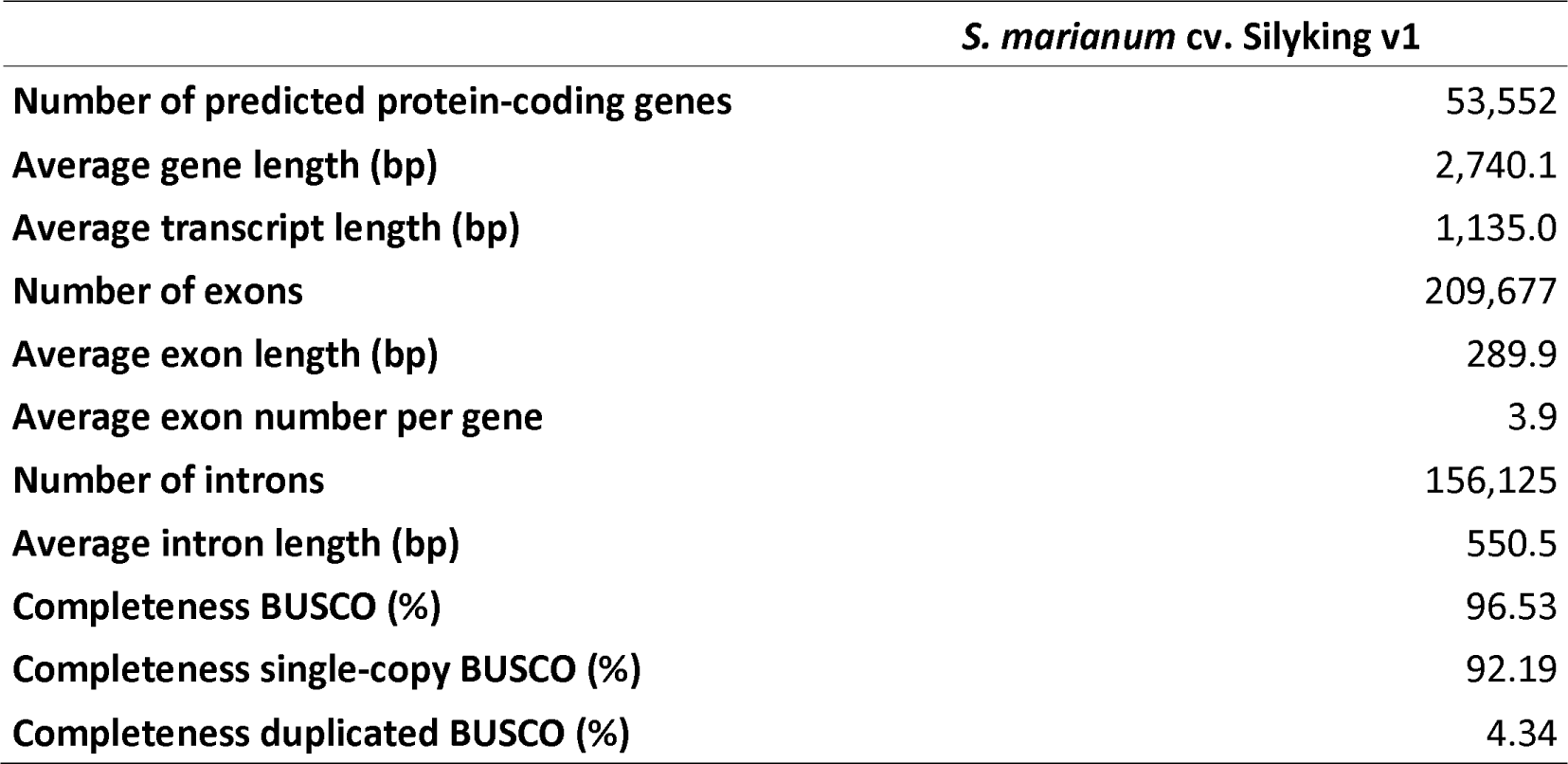
Summary of gene annotation of *Silybum marianum* cv. Silyking v1.

### Comparative genomic analysis

Collinearity in the *S. marianum* genome was identified through MCScanX^42^ and visualized with Circos v.0.66^43^. Additionally, chromosomal level collinearity was assessed between *S. marianum*, *Cynara cardunculus* , and *Helianthus annuus* using MCScanX^42^ and PanSyn v1.0. The collinearity between *S. marianum* and *C. cardunculus* was highly conserved showing a 1-to-1 relationship of chromosomes (Figure 4A and 4B), while that between *S. marianum* and *H. annuus* showed complex and discontiguous patterns (Figure 4C and 4D).

**Figure 4.**
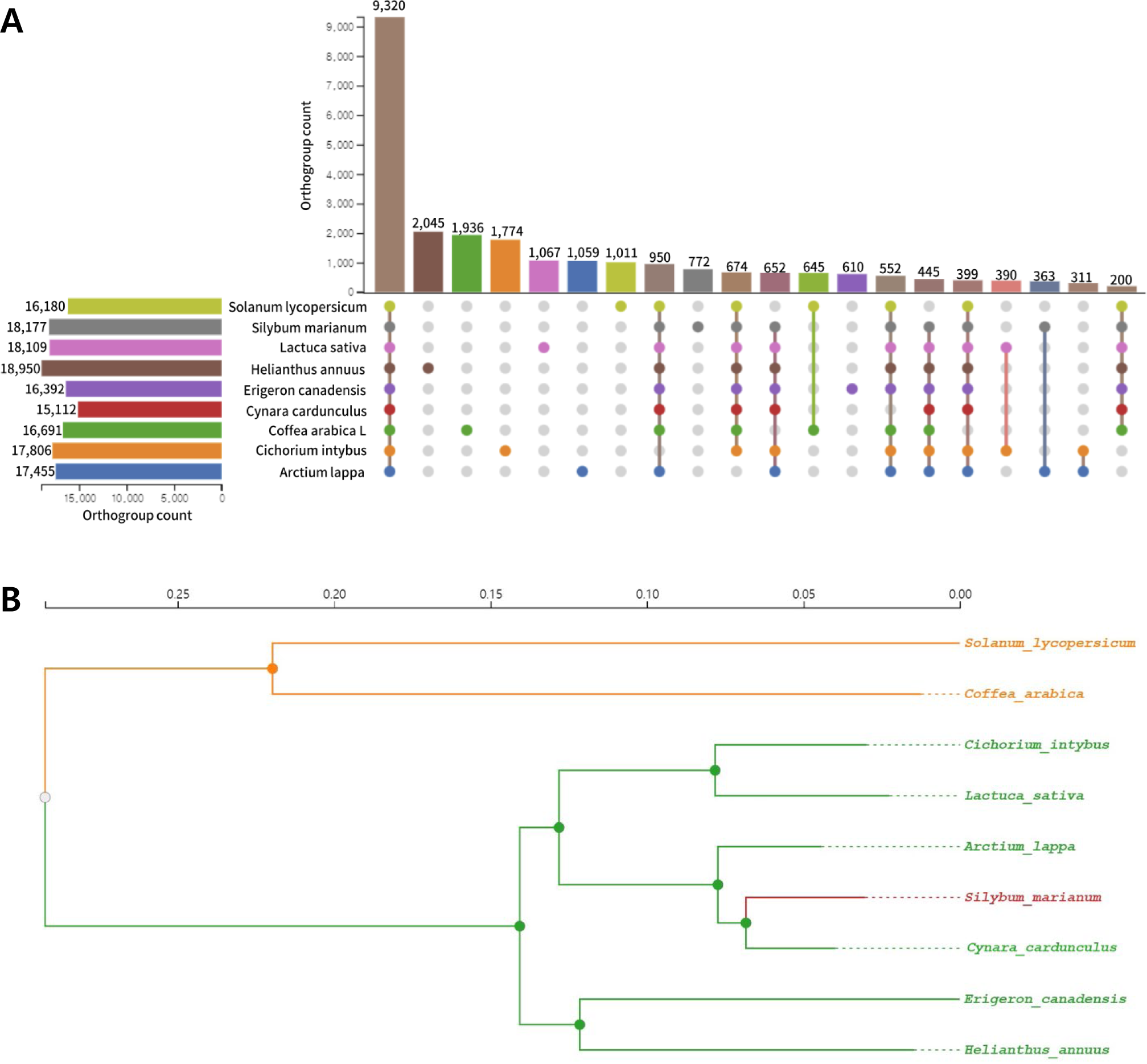
Genome evolution of *Silybum marianum* . (A) Top 20 orthogroups between *S. marianum* and eight plant species. See Supplementary Table 5 for the number of genes per orthogroup. (B) Phylogenetic tree of *S. marianum* and eight plant species.

By using OrthoFinder2 v2.3.12^44^ and the protein sequences, orthogroups were identified between *S. marianum* and eight species (Supplementary Table 4), including *C. cardunculus* (Artichoke, GCA_001531365.2)^48^, *H. annuus* (Common sunflower, GCA_002127325.1)^49^*, Arctium lappa* (Great burdock, GCA_023525745.1)^50^*, Cichorium intybus* (Chicory, GCA_023525715.1)^51^, *Erigeron canadensis* (Horseweed, GCA_010389155.1)^52^, *Lactuca sativa* (Lettuce, GCA_002870075.3)^53^, *Solanum lycopersicum* (Tomato, ITAG4.0), and *Coffea Arabica L*. (Coffee, GCA_003713225.1)^54^. A total of 31,351 orthogroups were identified, comprising 263,955 genes in total (Figure 5A, Supplementary Table 5). The phylogenetic tree was constructed using FastTree2^45^ based on the multiple sequence alignments of clustered orthogroups performed using MAFFT v7.3.13^29^ (Figure 5B).

### Data Records

Chromosome-level genome assembly of *S. marianum* has been deposited at the NCBI GenBank under accession number JAWIMA000000000^55^. Raw data for nanopore sequencing and RNA-seq have been deposited at the NCBI Sequence Read Archive under accession numbers SRR28145636-SRR28145644^56–62^, and are currently available under accession number PRJNA1021369 (https://www.ncbi.nlm.nih.gov/bioproject/PRJNA1021369). The sequences of genome assembly, the annotations of genes and transposable elements, and the list of orthogroups between *S. marianum* and eight species are available at Figshare^46^.

### Technical Validation

#### Genome assembly and gene prediction

For the quality assessment of genome assembly, we aligned the sequence reads from both RNA-seq and whole-genome resequencing data into our assembly, showing 97.6% and 99.4% of trimmed reads aligned, respectively (Table 1). Additionally, we checked the completeness of our assembly using BUSCO v4.1.4 with the embryophyte_odb10 database. As a result, the genome assembly from the previous research (ASM154182v1) showed 36.7% of completeness while our assembly from this study showed 99.1% of completeness. Moreover, the continuity of our assembly was evaluated using the LTR Assembly Index (LAI)^47^. The LAI score of our assembly was 17.77, which was higher than that of the Arabidopsis reference genome (TAIR10; LAI = 14.9). Our genome assembly can be considered as ‘reference quality’ with an LAI score ranging from 10 to 20, proposed by Ou et al. (2018)^47^.

For the validation of gene prediction, we used BUSCO with embryophyte_odb10 and viridiplantae_odb10 databases (Supplementary Table 6). With the embryophyte database, the predicted *S. marianum* protein-coding genes showed 96.53% of completeness. In the case of the viridiplantae database, predicted *S. marianum* protein-coding genes showed 97.41% of completeness.

#### Functional annotation of protein-coding genes

Functional annotation of the predicted genes identified 53,552 genes in *S. marianum* (Supplementary Table 7). More than 97% (51,994 genes) of predicted genes showed homology with the sequences in the NCBI RefSeq database. Moreover, 50,329 genes (94% of total genes) with functional descriptions in public databases such as NCBI RefSeq, Uniprot, and TAIR were categorized as known proteins. Additionally, 1,853 genes aligned by BLAST but lacking a characterized term and 1,370 genes not aligned by BLAST but showing FPKM>0.5 in RNA-Seq were categorized as uncharacterized genes.

## Code Availability

All data processed with publicly available bioinformatics tools or pipelines followed the analysis guidelines provided by those tools. No custom code was used during this study for the curation and/or validation of the dataset.

## Acknowledgements

This research was supported by the Rural Development Administration under the project No. PJ015988.

## Author contributions

K.K., S.H., and J.C. designed the study and led the research. J.S. collected and developed plant materials. K.L. performed genome assembly. M.E. performed TE annotation and LTR age estimation. J.H., D.K., and Y.Y. performed gene prediction, functional annotation, and comparative genomic analysis. B.A. and S.H. provided the research funding. K.K, J.H., and S.P. draft the manuscript. K.K. and J.C. revised the manuscript. All authors have read and agreed to the published version of the manuscript.

## Figures

provided as separate files

**Supplementary Figure 1.**
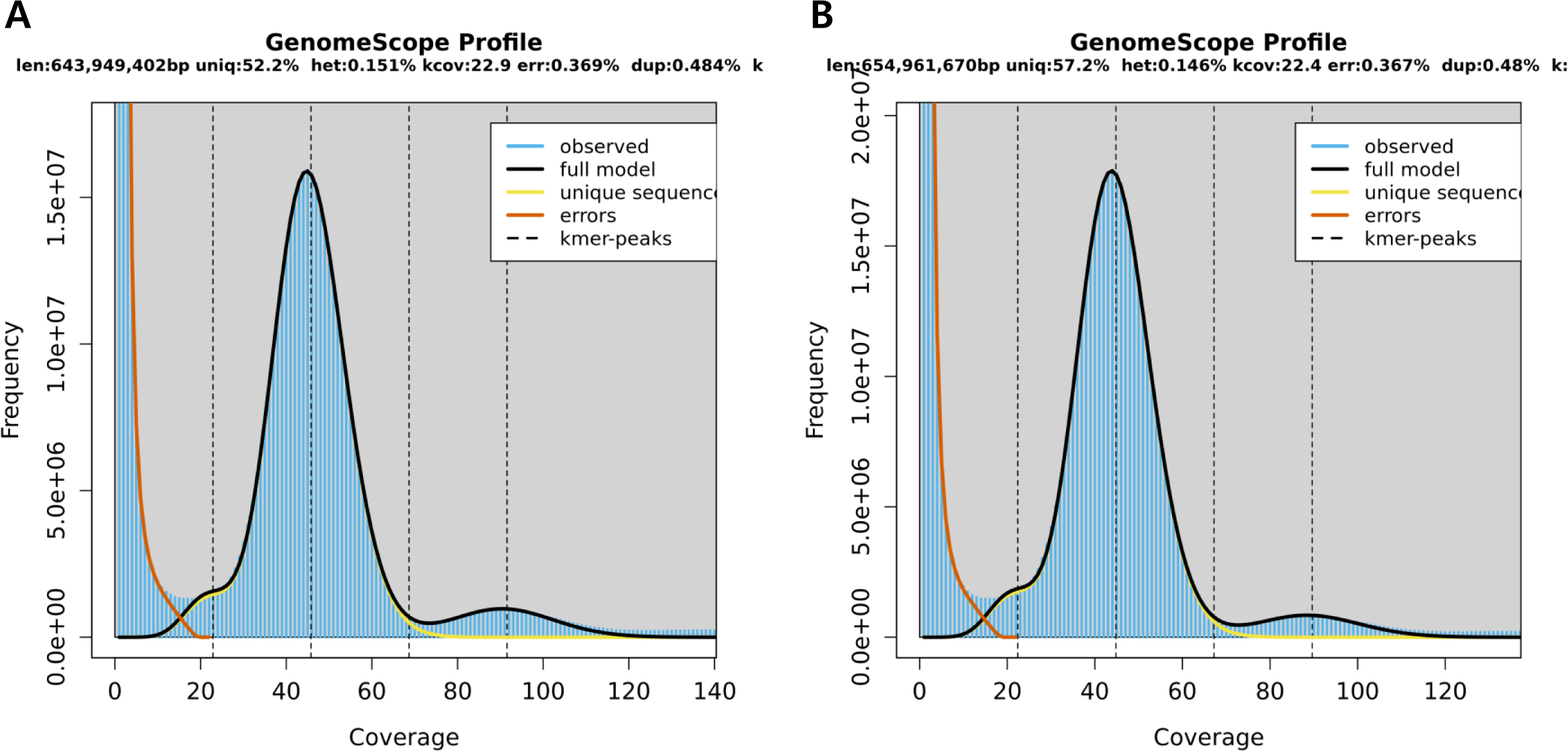
Genome survey of *Silybum marianum* using k-mer analysis. (A) GenomeScope profile using 19-mer. The genome size was estimated as 643 Mb with 0.14% heterozygosity. (B) GenomeScope profile using 21-mer. The genome size was estimated as 654 Mb with 0.14% heterozygosity.

**Supplementary Figure 2.**
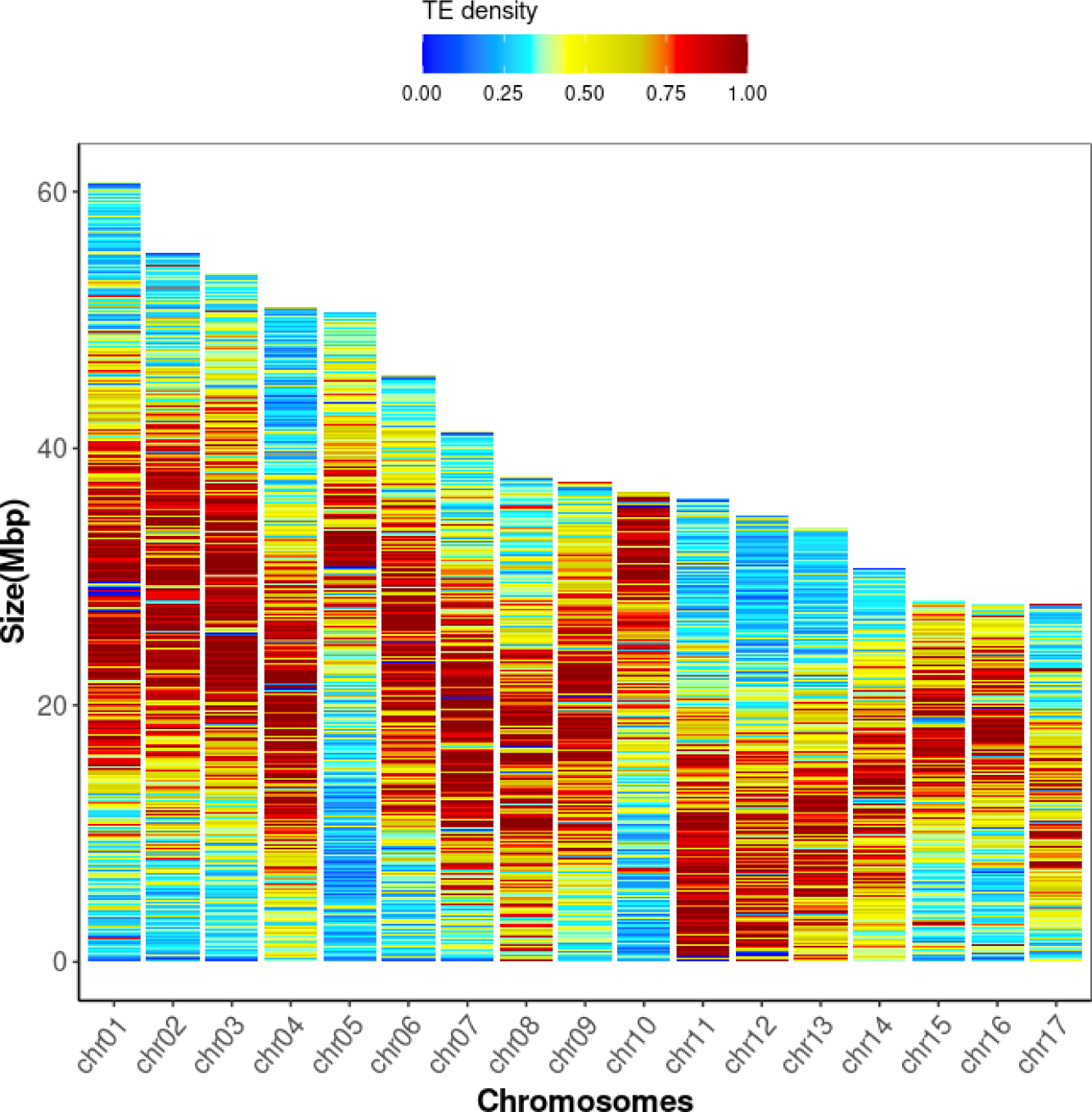
Heatmap of TE density across 17 chromosomes. The color intensity represents the level of TE density. Red color shows the high TE density while blue color shows the low TE density.

**Supplementary Figure 3.**
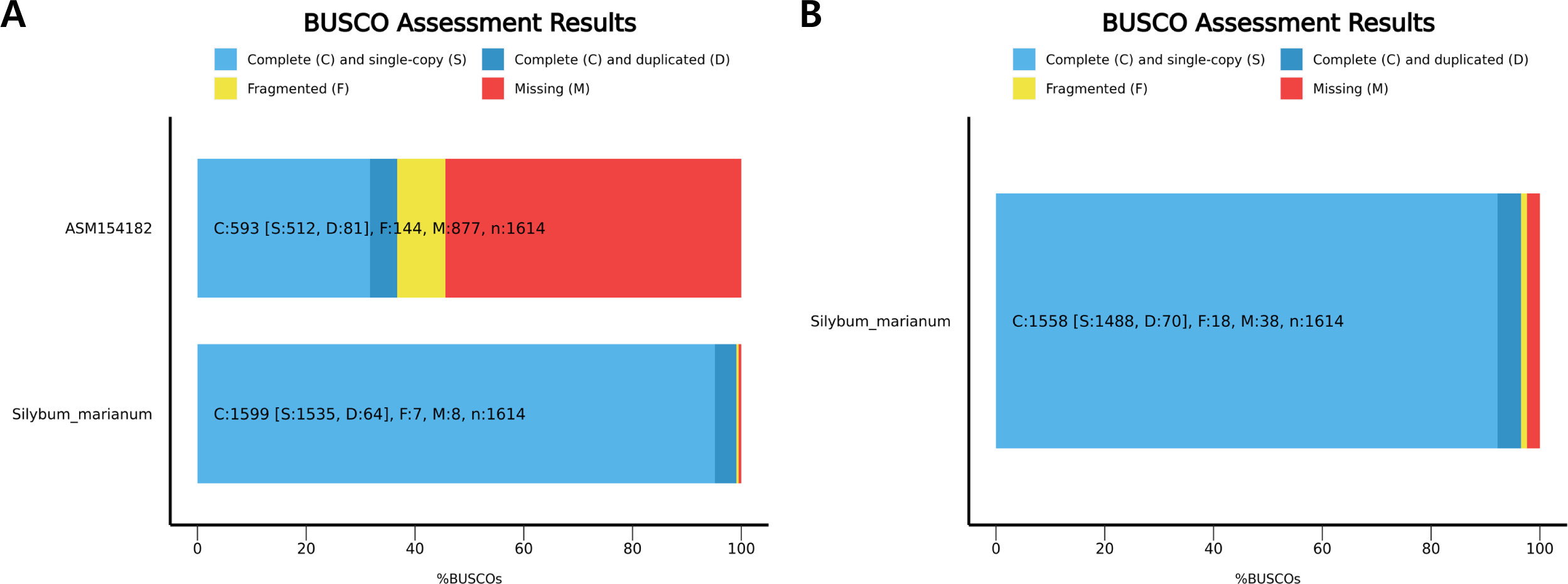
BUSCO assessment of *Silybum marianum* . A core gene set of embryophytes_odb10 was used for the assessment. (A) BUSCO completeness assessment for genome assemblies of *S. marianum*. Top: ASM154182v1, Bottom: cv. Silyking v1. (B) BUSCO completeness assessment for gene annotation of *S. marianum* cv. Silyking v1.

**Supplementary Table 1.**
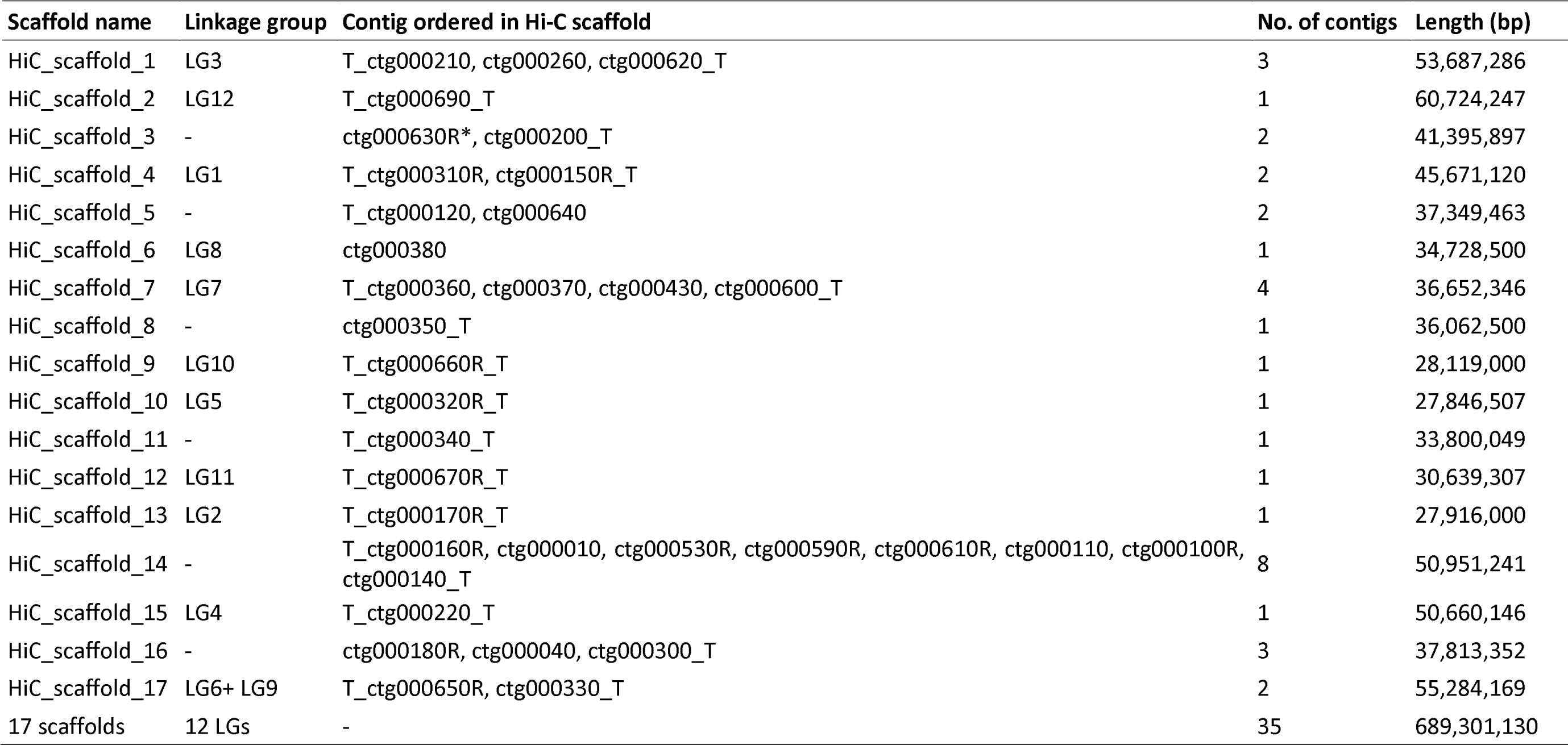
Summary of Pore-C scaffolding.

**Supplementary Table 2.**
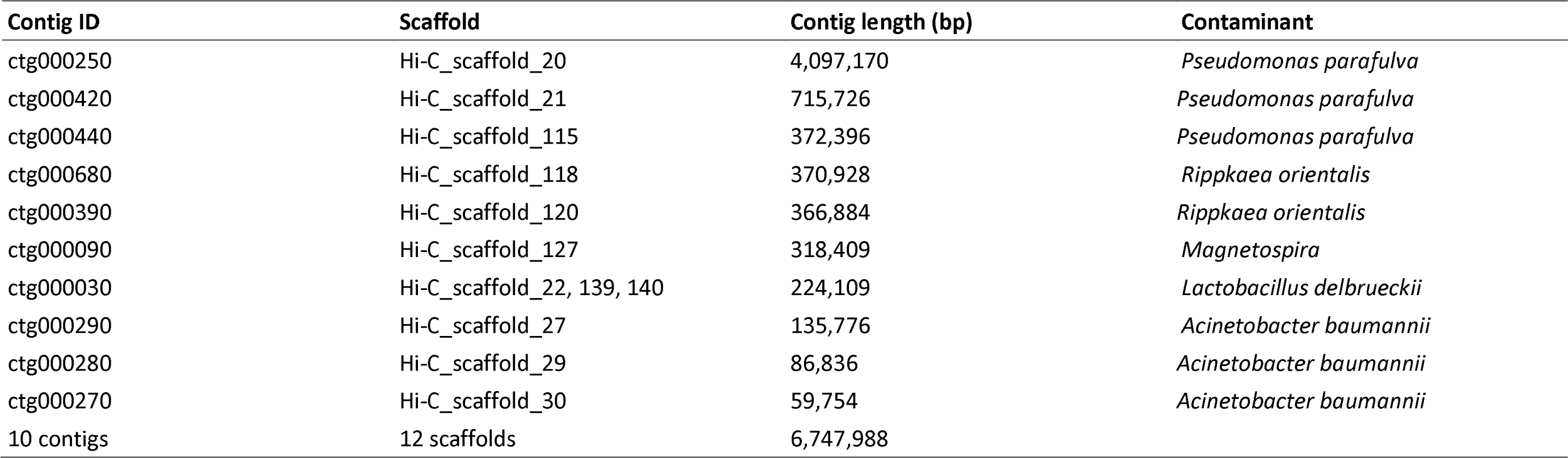
Unplaced contigs showing high similarity with bacterial sequences.

**Supplementary Table 3.**
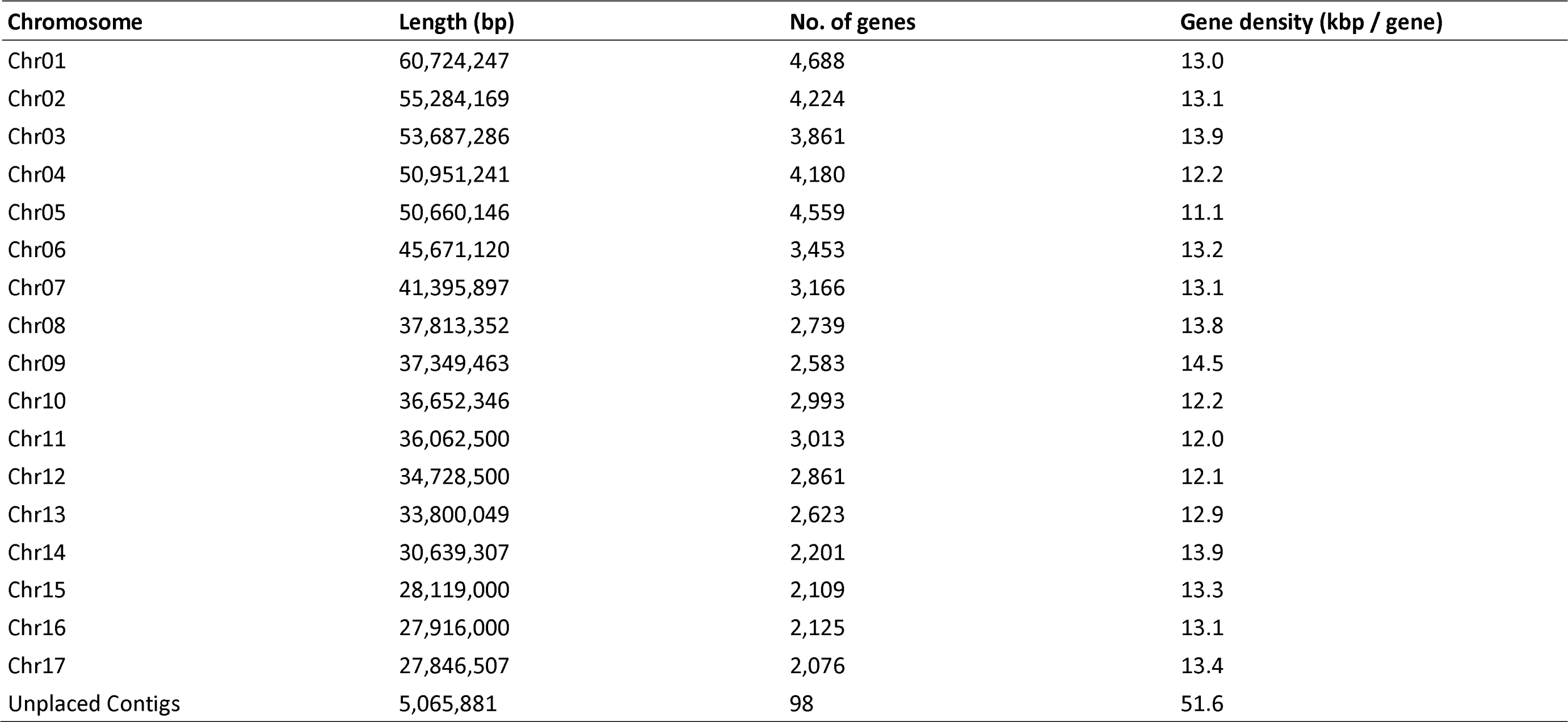
Chromosome-level summary of genome assembly and gene annotation of *Silybum marianum* cv. Silyking v1. some Length (bp) No. of genes Gene density (kbp / gene)

**Supplementary Table 4.**
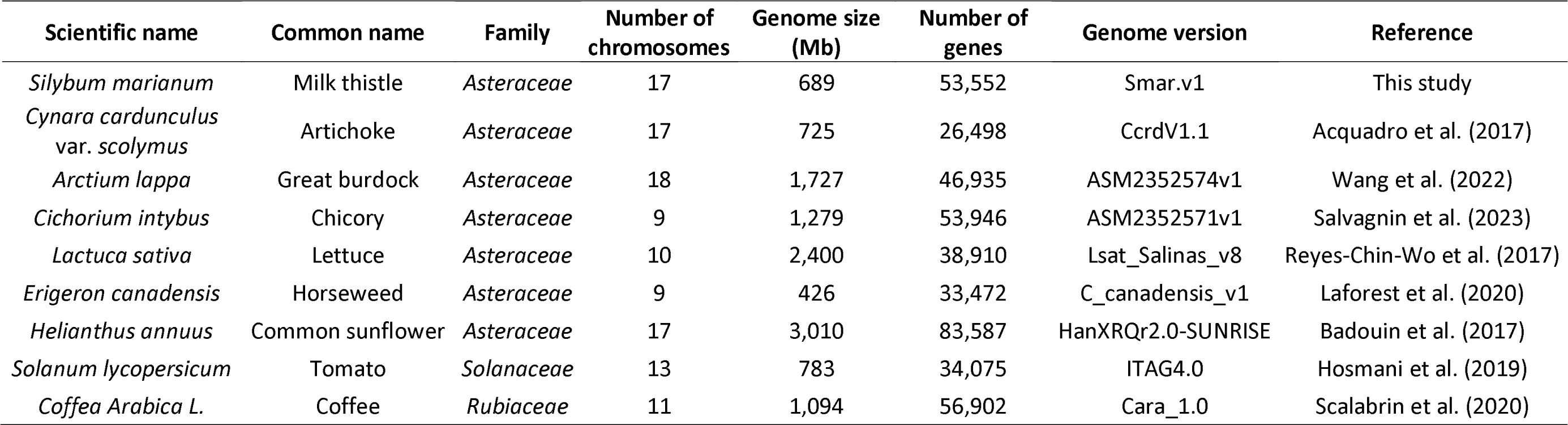
Summary of nine plant species for comparative genomic analysis in this study.

**Supplementary Table 5.**
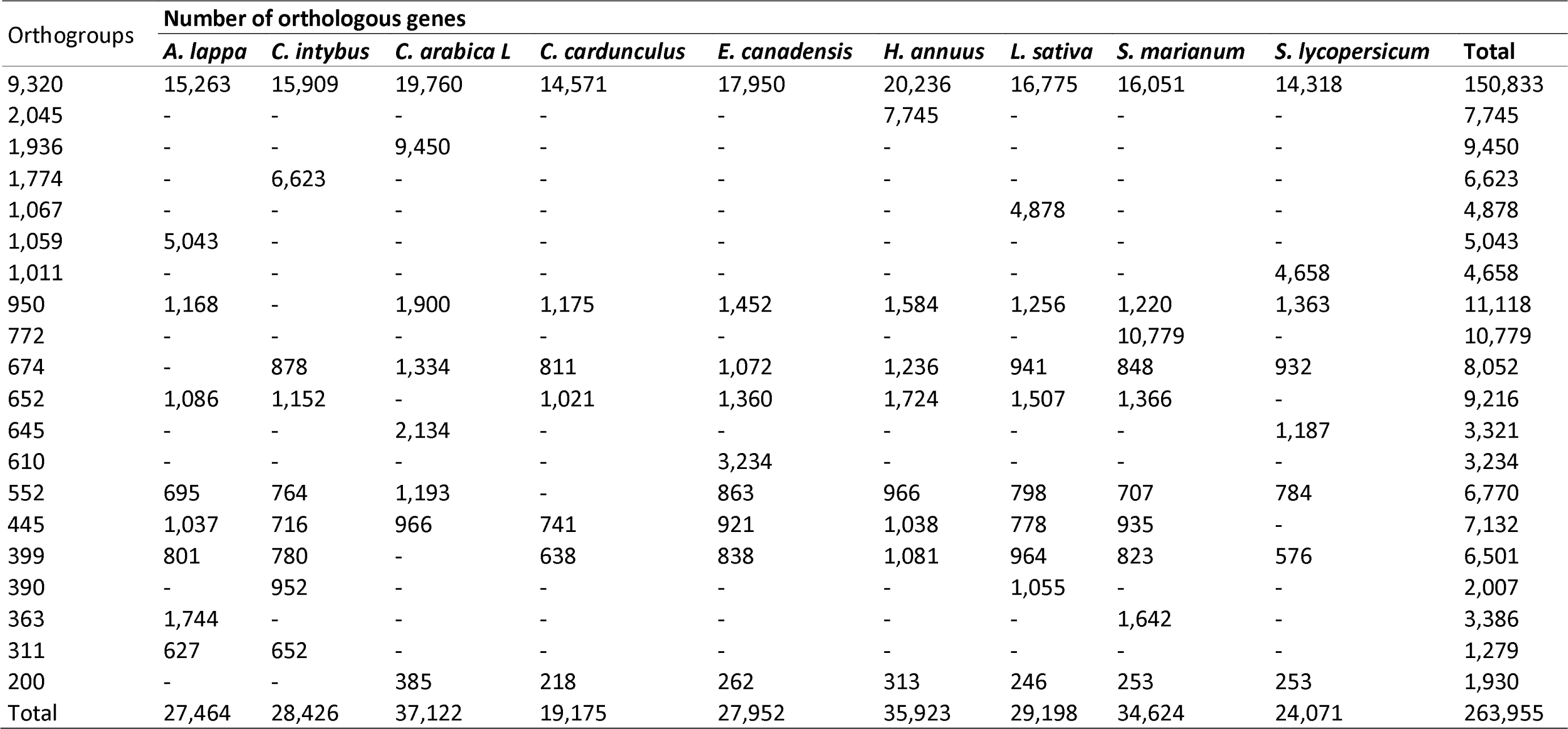
Orthologous genes from orthogroups determined across nine plant species.

**Supplementary Table 6.**
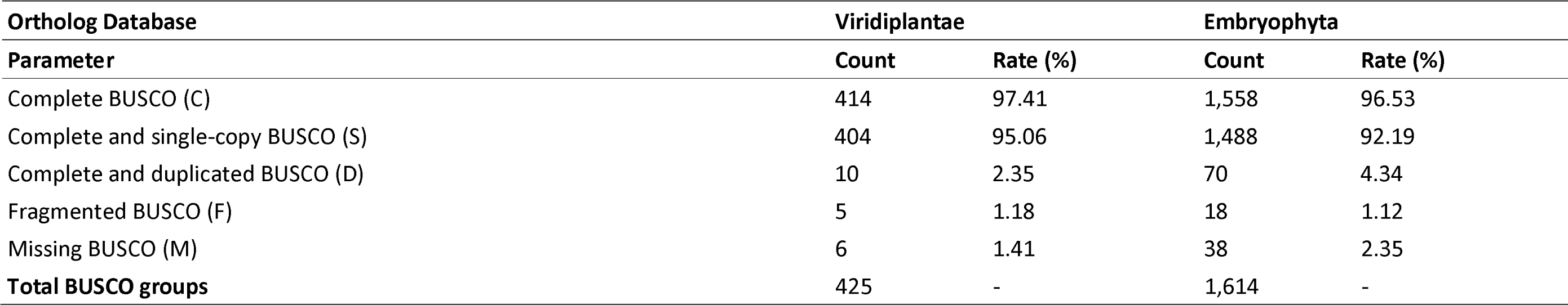
BUSCO assessment of gene annotation of *Silybum marianum* cv. Silyking v1.

**Supplementary Table 7.**
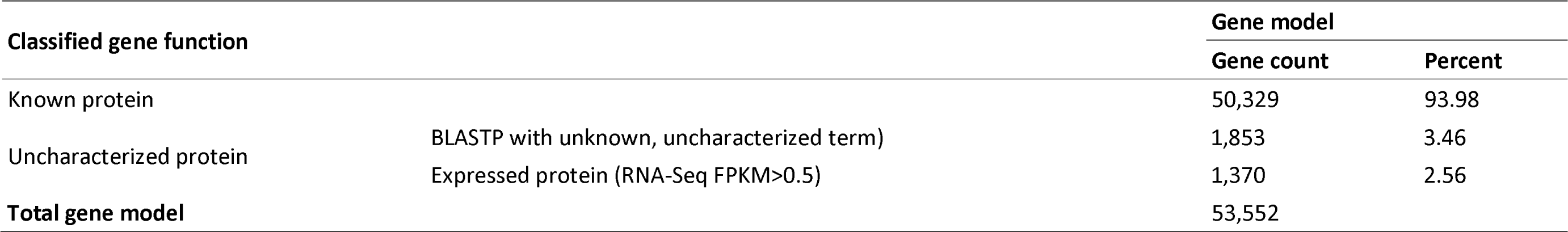
Summary of functional gene annotation of *Silybum marianum*cv. Silyking v1.

## References

1 Marceddu, R., Dinolfo, L., Carrubba, A., Sarno, M. & Di Miceli, G. Milk Thistle (Silybum Marianum L.) as a Novel Multipurpose Crop for Agriculture in Marginal Environments: A Review. Agronomy 12, 729 (2022).

2 Liava, V., Ntatsi, G. & Karkanis, A. Seed Germination of Three Milk Thistle (Silybum marianum (L.) Gaertn.) Populations of Greek Origin: Temperature, Duration, and Storage Conditions Effects. Plants (Basel) 12, doi:10.3390/plants12051025 (2023).

3 Young, J., Evans, R. & Hawkes, R. Milk thistle (Silybum marianum) seed germination. Weed Science 26, 395–398 (1978).

4 Corchete, P. in Bioactive molecules and medicinal plants 123–148 (Springer, 2008).

5 Schadewaldt, H. The history of Silymarin. Contribution to the history of liver therapy. Die Medizinische Welt 20, 902–914 (1969).

6 Lee, D. Y.-W. & Liu. Molecular structure and stereochemistry of silybin a, silybin B, isosilybin a, and isosilybin B, isolated from Silybum m arianum (milk thistle). Journal of natural products 66, 1171–1174 (2003).

7 Malekzadeh, M., Mirmazloum, S., Mortazavi, S., Panahi, M. & Angorani, H. Physicochemical properties and oil constituents of milk thistle (Silybum marianum Gaertn. cv. Budakalászi) under drought stress. Journal of Medicinal Plants Research 5, 1485–1488 (2011).

8 Abourashed, E. A., Mikell, J. R. & Khan, I. A. Bioconversion of silybin to phase I and II microbial metabolites with retained antioxidant activity. Bioorganic & medicinal chemistry 20, 2784–2788 (2012).

9 Polyak, S. J. *et a l* Identification of hepatoprotective flavonolignans from silymarin. Proceedings of the national academy of sciences 107, 5995–5999 (2010).

10 Saller, R., Brignoli, R., Melzer, J. & Meier, R. An updated systematic review with meta-analysis for the clinical evidence of silymarin. Forsch Komplementmed 15, 9–20, doi:10.1159/000113648 (2008).

11 Rainone, F. Milk thistle. American family physician 72, 1285–1292 (2005).

12 Flora, K., Hahn, M., Rosen, H. & Benner, K. Milk thistle (Silybum marianum) for the therapy of liver disease. Am J Gastroenterol 93, 139–143, doi:10.1111/j.1572-0241.1998.00139.x (1998).

13 Saller, R., Meier, R. & Brignoli, R. The use of silymarin in the treatment of liver diseases. Drugs 61, 2035–2063 (2001).

14 Vargas-Mendoza, N. et al. Hepatoprotective effect of silymarin. World journal of hepatology 6, 144 (2014).

15 Deep, G., Oberlies, N. H., Kroll, D. J. & Agarwal, R. Identifying the differential effects of silymarin constituents on cell growth and cell cycle regulatory molecules in human prostate cancer cells. International journal of cancer 123, 41–50 (2008).

16 Toyang, N. J. & Verpoorte, R. A review of the medicinal potentials of plants of the genus Vernonia (Asteraceae). Journal of Ethnopharmacology 146, 681–723 (2013).

17 Abenavoli, L., Capasso, R., Milic, N. & Capasso, F. Milk thistle in liver diseases: past, present, future. Phytotherapy Research 24, 1423–1432 (2010).

18 Bhattacharya, S. Phytotherapeutic properties of milk thistle seeds: An overview. J Adv Pharm Educ Res 1, 69–79 (2011).

19 Valková, V., Ďúranová, H., Bilčíková, J. & Habán, M. Milk thistle (Silybum marianum): a valuable medicinal plant with several therapeutic purposes. *The Journal of Microbiology*, Biotechnology and Food Sciences 9, 836 (2020).

20 Bolger, A. M., Lohse, M. & Usadel, B. Trimmomatic: a flexible trimmer for Illumina sequence data. Bioinformatics 30, 2114–2120, doi:10.1093/bioinformatics/btu170 (2014).

21 Marcais, G. & Kingsford, C. A fast, lock-free approach for efficient parallel counting of occurrences of k-mers. Bioinformatics 27, 764–770, doi:10.1093/bioinformatics/btr011 (2011).

22 Vurture, G. W. et al. GenomeScope: fast reference-free genome profiling from short reads. Bioinformatics 33, 2202–2204, doi:10.1093/bioinformatics/btx153 (2017).

23 Hu, J. et al. An efficient error correction and accurate assembly tool for noisy long reads. bioRxiv, 2023.2003.2009.531669, doi:10.1101/2023.03.09.531669 (2023).

24 Hu, J., Fan, J., Sun, Z. & Liu, S. NextPolish: a fast and efficient genome polishing tool for long-read assembly. Bioinformatics 36, 2253–2255, doi:10.1093/bioinformatics/btz891 (2019).

25 Durand, N. C., *et al .* Juicebox Provides a Visualization System for Hi-C Contact Maps with Unlimited Zoom. Cell Systems 3, 99–101, 10.1016/j.cels.2015.07.012 (2016).

26 Neumann, P., Novak, P., Hostakova, N. & Macas, J. Systematic survey of plant LTR-retrotransposons elucidates phylogenetic relationships of their polyprotein domains and provides a reference for element classification. Mob DNA 10, 1, doi:10.1186/s13100-018-0144-1 (2019).

27 Miele, V., Penel, S. & Duret, L. Ultra-fast sequence clustering from similarity networks with SiLiX. BMC Bioinformatics 12, 116, doi:10.1186/1471-2105-12-116 (2011).

28 Wang, M. & Kong, L. pblat: a multithread blat algorithm speeding up aligning sequences to genomes. BMC Bioinformatics 20, 28, doi:10.1186/s12859-019-2597-8 (2019).

29 Katoh, K. & Standley, D. M. MAFFT Multiple Sequence Alignment Software Version 7: Improvements in Performance and Usability. Molecular Biology and Evolution 30, 772–780, doi:10.1093/molbev/mst010 (2013).

30 Ellinghaus, D., Kurtz, S. & Willhoeft, U. LTRharvest, an efficient and flexible software for de novo detection of LTR retrotransposons. BMC Bioinformatics 9, 18, doi:10.1186/1471-2105-9-18 (2008).

31 Siren, J., Valimaki, N. & Makinen, V. Indexing Graphs for Path Queries with Applications in Genome Research. IEEE/ACM Trans Comput Biol Bioinform 11, 375–388, doi:10.1109/TCBB.2013.2297101 (2014).

32 Hoff, K. J., Lange, S., Lomsadze, A., Borodovsky, M. & Stanke, M. BRAKER1: Unsupervised RNA-Seq-Based Genome Annotation with GeneMark-ET and AUGUSTUS. Bioinformatics 32, 767–769, doi:10.1093/bioinformatics/btv661 (2015).

33 Lomsadze, A., Burns, P. D. & Borodovsky, M. Integration of mapped RNA-Seq reads into automatic training of eukaryotic gene finding algorithm. Nucleic Acids Res 42, e119, doi:10.1093/nar/gku557 (2014).

34 Ter-Hovhannisyan, V., Lomsadze, A., Chernoff, Y. O. & Borodovsky, M. Gene prediction in novel fungal genomes using an ab initio algorithm with unsupervised training. Genome Res 18, 1979–1990, doi:10.1101/gr.081612.108 (2008).

35 Stanke, M., Diekhans, M., Baertsch, R. & Haussler, D. Using native and syntenically mapped cDNA alignments to improve de novo gene finding. Bioinformatics 24, 637–644, doi:10.1093/bioinformatics/btn013 (2008).

36 Grabherr, M. G. et al. Full-length transcriptome assembly from RNA-Seq data without a reference genome. Nature Biotechnology 29, 644–652, doi:10.1038/nbt.1883 (2011).

37 Kovaka, S. et al. Transcriptome assembly from long-read RNA-seq alignments with StringTie2. Genome Biology 20, 278, doi:10.1186/s13059-019-1910-1 (2019).

38 Haas, B. J. et al. Improving the Arabidopsis genome annotation using maximal transcript alignment assemblies. Nucleic Acids Research 31, 5654–5666, doi:10.1093/nar/gkg770 (2003).

39 Slater, G. S. C. & Birney, E. Automated generation of heuristics for biological sequence comparison. BMC Bioinformatics 6, 31 (2005).

40 Haas, B. J. et al. Automated eukaryotic gene structure annotation using EVidenceModeler and the Program to Assemble Spliced Alignments. Genome Biol 9, R7, doi:10.1186/gb-2008-9-1-r7 (2008).

41 Jones, P. et al. InterProScan 5: genome-scale protein function classification. Bioinformatics 30, 1236–1240, doi:10.1093/bioinformatics/btu031 (2014).

42 Wang, Y. et al. MCScanX: a toolkit for detection and evolutionary analysis of gene synteny and collinearity. Nucleic Acids Research 40, e49–e49, doi:10.1093/nar/gkr1293 (2012).

43 Krzywinski, M. et al. C.ircos: an information aesthetic for comparative genomics. Genome Res 19, 1639–1645, doi:10.1101/gr.092759.109 (2009).

44 Emms, D. M. & Kelly, S. OrthoFinder: phylogenetic orthology inference for comparative genomics. Genome Biology 20, 238, doi:10.1186/s13059-019-1832-y (2019).

45 Price, M. N., Dehal, P. S. & Arkin, A. P. FastTree 2 – Approximately Maximum-Likelihood Trees for Large Alignments. PLOS ONE 5, e9490, doi:10.1371/journal.pone.0009490 (2010).

46 Kim, K. D. *Silybum marianum* genome assembly and annotation. Figshare 10.6084/m9.figshare.24190023.v2 (2024).

47 Ou, S., Chen, J. & Jiang, N. Assessing genome assembly quality using the LTR Assembly Index (LAI). Nucleic Acids Res 46, e126, doi:10.1093/nar/gky730 (2018).

48 NCBI GenBank https://identifiers.org/ncbi/insdc.gca:GCA_001531365.2 (2018).

49 NCBI GenBank https://identifiers.org/ncbi/insdc.gca:GCA_002127325.1 (2017).

50 NCBI GenBank https://identifiers.org/ncbi/insdc.gca:GCA_023525745.1 (2022).

51 NCBI GenBank https://identifiers.org/ncbi/insdc.gca:GCA_023525715.1 (2022).

52 NCBI GenBank https://identifiers.org/ncbi/insdc.gca:GCA_010389155.1 (2020).

53 NCBI GenBank https://identifiers.org/ncbi/insdc.gca:GCA_002870075.3 (2020).

54 NCBI GenBank https://identifiers.org/ncbi/insdc.gca:GCA_003713225.1 (2018).

55 NCBI GenBank https://identifiers.org/ncbi/insdc:JAWIMA000000000 (2024).

56 NCBI Sequence Read Archive https://identifiers.org/ncbi/insdc.sra:SRR28145636 (2024).

57 NCBI Sequence Read Archive https://identifiers.org/ncbi/insdc.sra:SRR28145637 (2024).

58 NCBI Sequence Read Archive https://identifiers.org/ncbi/insdc.sra:SRR28145638 (2024).

59 NCBI Sequence Read Archive https://identifiers.org/ncbi/insdc.sra:SRR28145639 (2024).

60 NCBI Sequence Read Archive https://identifiers.org/ncbi/insdc.sra:SRR28145640 (2024).

61 NCBI Sequence Read Archive https://identifiers.org/ncbi/insdc.sra:SRR28145641 (2024).

62 NCBI Sequence Read Archive https://identifiers.org/ncbi/insdc.sra:SRR28145642 (2024).

