## Supplementary material for "Chromosome-level genome assembly of milk thistle (*Silybum marianum* (L.) Gaertn.)": Tables

**Table 1. Comparison of genome assemblies between *Silybum marianum* ASM154182v1 and cv. Silyking v1.**

|  | | ***S. marianum* ASM154182v1** | ***S. marianum* cv. Silyking v1 (this study)** |
| --- | --- | --- | --- |
| **Long-read sequencing platform** | | PacBio | Oxford Nanopore |
| **Genome coverage** | Long-read (fold) | - | 111.3 |
|  | Short-read (fold) | - | 75.23 |
|  | Hi-C (bp; Total / N50) | - | 39.5 Gb / 5.4 kb |
| **Estimated genome size (Mb)** | | 1,477.58 | 694.4 |
| **Estimated heterozygosity (%)** | | - | 0.172 |
| **Number of scaffolds** | | - | 17 |
| **Total length of scaffolds (Mb)** | | - | 689.3 |
| **Scaffold N50 (Mb)** | | - | 41.4 |
| **Longest scaffold (Mb)** | | - | 60.7 |
| **Number of contigs** | | 258,575 | 70 |
| **Total length of contigs (Mb)** | | 1,477.57 | 5.1 |
| **Contig N50 (Mb)** | | 0.006967 | 0.25 |
| **Contig L50** | | 62,112 | 9 |
| **Longest contig (Mb)** | | 0.099455 | 0.39 |
| **GC content (%)** | | 37.2 | 34.5 |
| **Mapping rate of Illumina reads (%)** | | - | 99.4 |
| **Completeness BUSCO (%)** | | 36.7 | 99.1 |
| **Completeness single-copy BUSCO (%)** | | 31.7 | 95.1 |
| **Completeness duplicated BUSCO (%)** | | 5.0 | 4.0 |

**Table 2. Summary of transposable elements in *Silybum marianum* cv. Silyking v1.**

| **Code** | **Class** | **Order** | **Superfamily** | **Total length (bp)** | **% of genome** | **% of TE** |
| --- | --- | --- | --- | --- | --- | --- |
| RLC | Class I | LTR | Copia | 165,634,194 | 23.85 | 41.15 |
| RLG | Class I | LTR | Gypsy | 73,223,946 | 10.55 | 18.18 |
| RLX | Class I | LTR | Unknown | 44,826,005 | 6.46 | 11.13 |
| RIL | Class I | LINEs | Unknown | 5,467,739 | 0.79 | 1.36 |
| DTC | Class II | TIR | CACTA | 12,834,335 | 1.85 | 3.19 |
| DTM | Class II | TIR | Mutator | 14,770,974 | 2.13 | 3.67 |
| DTA | Class II | TIR | hAT | 2,264,852 | 0.33 | 0.56 |
| DXX | Class II | TIR | MITEs | 77,739,850 | 11.20 | 19.31 |
| DHH | Class II | Helitron | Helitron | 5,879,725 | 0.85 | 1.46 |
| Total |  |  |  | 402,641,620 | 58.00 % |  |

**Table 3. Summary of gene annotation of *Silybum marianum* cv. Silyking v1.**

|  | ***S. marianum* cv. Silyking v1** |
| --- | --- |
| **Number of predicted protein-coding genes** | 53,552 |
| **Average gene length (bp)** | 2,740.1 |
| **Average transcript length (bp)** | 1,135.0 |
| **Number of exons** | 209,677 |
| **Average exon length (bp)** | 289.9 |
| **Average exon number per gene** | 3.9 |
| **Number of introns** | 156,125 |
| **Average intron length (bp)** | 550.5 |
| **Completeness BUSCO (%)** | 96.53 |
| **Completeness single-copy BUSCO (%)** | 92.19 |
| **Completeness duplicated BUSCO (%)** | 4.34 |

**Supplementary Table 1. Summary of Pore-C scaffolding.**

| **Scaffold name** | **Linkage group** | **Contig ordered in Hi-C scaffold** | **No. of contigs** | **Length (bp)** |
| --- | --- | --- | --- | --- |
| HiC_scaffold_1 | LG3 | T_ctg000210, ctg000260, ctg000620_T | 3 | 53,687,286 |
| HiC_scaffold_2 | LG12 | T_ctg000690_T | 1 | 60,724,247 |
| HiC_scaffold_3 | - | ctg000630R*, ctg000200_T | 2 | 41,395,897 |
| HiC_scaffold_4 | LG1 | T_ctg000310R, ctg000150R_T | 2 | 45,671,120 |
| HiC_scaffold_5 | - | T_ctg000120, ctg000640 | 2 | 37,349,463 |
| HiC_scaffold_6 | LG8 | ctg000380 | 1 | 34,728,500 |
| HiC_scaffold_7 | LG7 | T_ctg000360, ctg000370, ctg000430, ctg000600_T | 4 | 36,652,346 |
| HiC_scaffold_8 | - | ctg000350_T | 1 | 36,062,500 |
| HiC_scaffold_9 | LG10 | T_ctg000660R_T | 1 | 28,119,000 |
| HiC_scaffold_10 | LG5 | T_ctg000320R_T | 1 | 27,846,507 |
| HiC_scaffold_11 | - | T_ctg000340_T | 1 | 33,800,049 |
| HiC_scaffold_12 | LG11 | T_ctg000670R_T | 1 | 30,639,307 |
| HiC_scaffold_13 | LG2 | T_ctg000170R_T | 1 | 27,916,000 |
| HiC_scaffold_14 | - | T_ctg000160R, ctg000010, ctg000530R, ctg000590R, ctg000610R, ctg000110, ctg000100R, ctg000140_T | 8 | 50,951,241 |
| HiC_scaffold_15 | LG4 | T_ctg000220_T | 1 | 50,660,146 |
| HiC_scaffold_16 | - | ctg000180R, ctg000040, ctg000300_T | 3 | 37,813,352 |
| HiC_scaffold_17 | LG6+ LG9 | T_ctg000650R, ctg000330_T | 2 | 55,284,169 |
| 17 scaffolds | 12 LGs | - | 35 | 689,301,130 |

**Supplementary Table 2. Unplaced contigs showing high similarity with bacterial sequences.**

| **Contig ID** | **Scaffold** | **Contig length (bp)** | **Contaminant** |
| --- | --- | --- | --- |
| ctg000250 | Hi-C_scaffold_20 | 4,097,170 | *Pseudomonas parafulva* |
| ctg000420 | Hi-C_scaffold_21 | 715,726 | *Pseudomonas parafulva* |
| ctg000440 | Hi-C_scaffold_115 | 372,396 | *Pseudomonas parafulva* |
| ctg000680 | Hi-C_scaffold_118 | 370,928 | *Rippkaea orientalis* |
| ctg000390 | Hi-C_scaffold_120 | 366,884 | *Rippkaea orientalis* |
| ctg000090 | Hi-C_scaffold_127 | 318,409 | *Magnetospira* |
| ctg000030 | Hi-C_scaffold_22, 139, 140 | 224,109 | *Lactobacillus delbrueckii* |
| ctg000290 | Hi-C_scaffold_27 | 135,776 | *Acinetobacter baumannii* |
| ctg000280 | Hi-C_scaffold_29 | 86,836 | *Acinetobacter baumannii* |
| ctg000270 | Hi-C_scaffold_30 | 59,754 | *Acinetobacter baumannii* |
| 10 contigs | 12 scaffolds | 6,747,988 |  |

**Supplementary Table 3. Chromosome-level summary of genome assembly and gene annotation of *Silybum marianum* cv. Silyking v1.**

| **Chromosome** | **Length (bp)** | **No. of genes** | **Gene density (kbp / gene)** |
| --- | --- | --- | --- |
| Chr01 | 60,724,247 | 4,688 | 13.0 |
| Chr02 | 55,284,169 | 4,224 | 13.1 |
| Chr03 | 53,687,286 | 3,861 | 13.9 |
| Chr04 | 50,951,241 | 4,180 | 12.2 |
| Chr05 | 50,660,146 | 4,559 | 11.1 |
| Chr06 | 45,671,120 | 3,453 | 13.2 |
| Chr07 | 41,395,897 | 3,166 | 13.1 |
| Chr08 | 37,813,352 | 2,739 | 13.8 |
| Chr09 | 37,349,463 | 2,583 | 14.5 |
| Chr10 | 36,652,346 | 2,993 | 12.2 |
| Chr11 | 36,062,500 | 3,013 | 12.0 |
| Chr12 | 34,728,500 | 2,861 | 12.1 |
| Chr13 | 33,800,049 | 2,623 | 12.9 |
| Chr14 | 30,639,307 | 2,201 | 13.9 |
| Chr15 | 28,119,000 | 2,109 | 13.3 |
| Chr16 | 27,916,000 | 2,125 | 13.1 |
| Chr17 | 27,846,507 | 2,076 | 13.4 |
| Unplaced Contigs | 5,065,881 | 98 | 51.6 |

**Supplementary Table 4. Summary of nine plant species for comparative genomic analysis in this study.**

| **Scientific name** | **Common name** | **Family** | **Number of chromosomes** | **Genome size (Mb)** | **Number of genes** | **Genome version** | **Reference** |
| --- | --- | --- | --- | --- | --- | --- | --- |
| *Silybum marianum* | Milk thistle | *Asteraceae* | 17 | 689 | 53,552 | Smar.v1 | This study |
| *Cynara cardunculus* var. *scolymus* | Artichoke | *Asteraceae* | 17 | 725 | 26,498 | CcrdV1.1 | Acquadro et al. (2017) |
| *Arctium lappa* | Great burdock | *Asteraceae* | 18 | 1,727 | 46,935 | ASM2352574v1 | Wang et al. (2022) |
| *Cichorium intybus* | Chicory | *Asteraceae* | 9 | 1,279 | 53,946 | ASM2352571v1 | Salvagnin et al. (2023) |
| *Lactuca sativa* | Lettuce | *Asteraceae* | 10 | 2,400 | 38,910 | Lsat_Salinas_v8 | Reyes-Chin-Wo et al. (2017) |
| *Erigeron canadensis* | Horseweed | *Asteraceae* | 9 | 426 | 33,472 | C_canadensis_v1 | Laforest et al. (2020) |
| *Helianthus annuus* | Common sunflower | *Asteraceae* | 17 | 3,010 | 83,587 | HanXRQr2.0-SUNRISE | Badouin et al. (2017) |
| *Solanum lycopersicum* | Tomato | *Solanaceae* | 13 | 783 | 34,075 | ITAG4.0 | Hosmani et al. (2019) |
| *Coffea Arabica L.* | Coffee | *Rubiaceae* | 11 | 1,094 | 56,902 | Cara_1.0 | Scalabrin et al. (2020) |

**Supplementary Table 5. Orthologous genes from orthogroups determined across nine plant species.**

| Orthogroups | **Number of orthologous genes** | | | | | | | | | |
| --- | --- | --- | --- | --- | --- | --- | --- | --- | --- | --- |
|  | ***A. lappa*** | ***C. intybus*** | ***C. arabica L*** | ***C. cardunculus*** | ***E. canadensis*** | ***H. annuus*** | ***L. sativa*** | ***S. marianum*** | ***S. lycopersicum*** | **Total** |
| 9,320 | 15,263 | 15,909 | 19,760 | 14,571 | 17,950 | 20,236 | 16,775 | 16,051 | 14,318 | 150,833 |
| 2,045 | - | - | - | - | - | 7,745 | - | - | - | 7,745 |
| 1,936 | - | - | 9,450 | - | - | - | - | - | - | 9,450 |
| 1,774 | - | 6,623 | - | - | - | - | - | - | - | 6,623 |
| 1,067 | - | - | - | - | - | - | 4,878 | - | - | 4,878 |
| 1,059 | 5,043 | - | - | - | - | - | - | - | - | 5,043 |
| 1,011 | - | - | - | - | - | - | - | - | 4,658 | 4,658 |
| 950 | 1,168 | - | 1,900 | 1,175 | 1,452 | 1,584 | 1,256 | 1,220 | 1,363 | 11,118 |
| 772 | - | - | - | - | - | - | - | 10,779 | - | 10,779 |
| 674 | - | 878 | 1,334 | 811 | 1,072 | 1,236 | 941 | 848 | 932 | 8,052 |
| 652 | 1,086 | 1,152 | - | 1,021 | 1,360 | 1,724 | 1,507 | 1,366 | - | 9,216 |
| 645 | - | - | 2,134 | - | - | - | - | - | 1,187 | 3,321 |
| 610 | - | - | - | - | 3,234 | - | - | - | - | 3,234 |
| 552 | 695 | 764 | 1,193 | - | 863 | 966 | 798 | 707 | 784 | 6,770 |
| 445 | 1,037 | 716 | 966 | 741 | 921 | 1,038 | 778 | 935 | - | 7,132 |
| 399 | 801 | 780 | - | 638 | 838 | 1,081 | 964 | 823 | 576 | 6,501 |
| 390 | - | 952 | - | - | - | - | 1,055 | - | - | 2,007 |
| 363 | 1,744 | - | - | - | - | - | - | 1,642 | - | 3,386 |
| 311 | 627 | 652 | - | - | - | - | - | - | - | 1,279 |
| 200 | - | - | 385 | 218 | 262 | 313 | 246 | 253 | 253 | 1,930 |
| Total | 27,464 | 28,426 | 37,122 | 19,175 | 27,952 | 35,923 | 29,198 | 34,624 | 24,071 | 263,955 |

**Supplementary Table 6. BUSCO assessment of gene annotation of *Silybum marianum* cv. Silyking v1.**

| **Ortholog Database** | **Viridiplantae** | | **Embryophyta** | |
| --- | --- | --- | --- | --- |
| **Parameter** | **Count** | **Rate (%)** | **Count** | **Rate (%)** |
| Complete BUSCO (C) | 414 | 97.41 | 1,558 | 96.53 |
| Complete and single-copy BUSCO (S) | 404 | 95.06 | 1,488 | 92.19 |
| Complete and duplicated BUSCO (D) | 10 | 2.35 | 70 | 4.34 |
| Fragmented BUSCO (F) | 5 | 1.18 | 18 | 1.12 |
| Missing BUSCO (M) | 6 | 1.41 | 38 | 2.35 |
| **Total BUSCO groups** | 425 | - | 1,614 | - |

**Supplementary Table 7. Summary of functional gene annotation of *Silybum marianum* cv. Silyking v1.**

| **Classified gene function** | | **Gene model** | |
| --- | --- | --- | --- |
|  |  | **Gene count** | **Percent** |
| Known protein | | 50,329 | 93.98 |
| Uncharacterized protein | BLASTP with unknown, uncharacterized term) | 1,853 | 3.46 |
|  | Expressed protein (RNA-Seq FPKM>0.5) | 1,370 | 2.56 |
| **Total gene model** | | 53,552 |  |
